# Accumulation of prelamin A drives inflammation in the heart with implications for treatment of inherited and acquired cardiomyopathies

**DOI:** 10.1101/457044

**Authors:** Daniel Brayson, Andrea Frustaci, Romina Verardo, Cristina Chimenti, Matteo Antonio Russo, Robert Hayward, Sadia Munir Ahmad, Gema Vizcay-Barrena, Andrea Protti, Peter S. Zammit, Cristobal G. dos Remedios, Elisabeth Ehler, Ajay M. Shah, Catherine M. Shanahan

## Abstract

Cardiomyopathies are complex heart muscle diseases that can be inherited e.g. dilated cardiomyopathy resulting from LMNA gene mutations, or acquired, e.g. cardiomyopathy associated with HIV. In both cases the lamin A precursor, prelamin A, may play a central role: mutations in *LMNA* and certain HIV protease inhibitors acting via the enzyme ZMPSTE24 both inhibit prelamin A processing. Firstly, we show that myocardial prelamin A accumulation occurs in both these cardiomyopathies in patients. Secondly, we developed a novel mouse model of cardiac specific prelamin A accumulation which mimicked tissue and molecular features of HIV associated cardiomyopathy, including inflammation. These findings: (1) confirm a central pathological role of prelamin A common to genetic and acquired cardiomyopathies; (2) have implications for the management of HIV patients with cardiac disease in whom protease inhibitors with low/no binding to ZMPSTE24 may be preferred; and (3) suggest that targeting inflammation may be a useful treatment strategy for some forms of inherited cardiomyopathy.

## Introduction

Mutations in the *LMNA* gene are commonly implicated in dilated cardiomyopathy (DCM) phenotypes (1), accounting for approximately 6% of all cases (2). Investigation of *in vivo* mouse models harbouring *LMNA* mutations associated with clinical DCM have identified a number of mechanisms associated with disease (3). However, some questions remain unresolved, in particular, whether the lamin A precursor, prelamin A is involved in the pathogenesis of cardiomyopathies (4–7).

The *LMNA* gene produces two distinct proteins, lamin A and lamin C, which together with the B-type lamins form the nuclear lamina which sits adjacent to the inner nuclear membrane (INM) of the nuclear envelope (NE), on the nucleoplasmic side (8). The primary role of the lamina is to provide structural stability to the nuclear environment and to anchor heterochromatin, thereby facilitating appropriate gene expression and efficient DNA damage repair (9, 10). Additionally, the lamina forms part of the linker of nucleoskeleton to cytoskeleton (LINC) complex, which mediates physical communication with the cytoplasmic environment enabling rapid responses to physical cues, a process termed mechanotransduction (11).

To achieve lamin A maturation, its precursor prelamin A, requires step-wise proteolytic processing (12). After translation, addition of farnesyl and carboxymethyl groups to a CAAX motif in the C terminus occurs, followed by an upstream cleavage exclusively mediated by zinc metalloroteinase STE24 homologue, ZMPSTE24, to yield mature lamin A (13–16). Retention of this farnesylated C terminal domain by lamin A precursors is pathologic and mutations in both the *LMNA* and *ZMPSTE24* genes that cause this retention are implicated in premature ageing disorders, such as Hutchinson Gilford progeria syndrome (HGPS), as well as DCM. HGPS patients develop cardiomegaly, atrial enlargement and age-dependent diastolic and systolic dysfunction and left ventricle (LV) hypertrophy (17–19), while the DCM causing mutation *LMNA*-R89L has been shown to result in aberrant processing and accumulation of prelamin A (5, 7). Moreover, a mutation in *LMNA* postulated to inhibit prelamin A processing which causes Dunnigan-type familial lipodystrophy, is also associated with cardiac dysfunction, with patients homozygous for this mutation having worse LV function indicating a dose-dependent effect (4). Additionally a mutation in *ZMPSTE24* known to confer a reduction in enzyme activity was found in a patient with metabolic syndrome and cardiomyopathy (6).

Another cause of prelamin A accumulation is via the pharmacological inhibition of ZMPSTE24 activity by Human Immunodeficiency Virus Protease Inhibitors (HIV PIs) used in the treatment of HIV. HIV PIs result in prelamin A accumulation in cells and potentially contribute to adverse effects (20). HIV patients have double the risk for developing cardiovascular disease than non-carriers (21). Moreover, HIV patients can develop HIV-associated cardiomyopathy (22), though the aetiology is complex (23). Previous studies have identified nucleoside reverse transcriptase inhibitors (NRTIs) used in conjunction with HIV PIs as responsible for the development of cardiomyopathy in HIV patients (24, 25). Presently, there is limited knowledge on the impact of HIV PIs on the development of cardiomyopathy though there is an attempt to ‘characterise heart function on antiretroviral therapy’ with the introduction of the CHART study (26). These points considered, we sought to investigate the extent and effects of prelamin A accumulation in the setting of cardiomyopathy.

## Results

### Prelamin A accumulates in HIV associated cardiomyopathy and DCM samples

Western blotting of myocardial biopsy samples from HIV+ patients being treated with highly active antiretroviral therapy and presenting with cardiomyopathy (Table 1) showed that prelamin A was abundant in HIV+ patient myocardium but not in a selection of DCM samples and non-failing controls (Fig. 1A). Immunohistochemistry showed focal expression of prelamin A in nuclei of cardiomyocyte (CM) and non-CM populations within the hearts of HIV+ patients with a number of CM nuclei showing highly aberrant morphologies (Fig. 1B). This was confirmed by analysis of the myocardial ultrastructure using electron microscopy which showed evidence of nuclear morphology defects and changes in the spatial organisation of heterochromatin in HIV+ myocardium (Fig. 1C,D). Immunofluorescence staining for prelamin A was performed on human DCM patient left ventricle (LV) samples (Supp. Table 3) and non-failing control samples (Supp. Table 4), and quantified (Fig. 1E). Sporadic and focal expression of prelamin A was observable in CM nuclei in both non-failing and DCM samples (Fig. 1F). However, in one DCM sample (DCM05) consistent CM nuclear rim staining was found in 71.5% of total CM nuclei.

**Table 1.** Clinical characteristics and antiretroviral treatment regime of five patients with HIV-associated cardiomyopathy

| HIV+ Patient ID | Age | Sex | Time under retroviral therapy, years | Echocardiography Data |  |
| --- | --- | --- | --- | --- | --- |
|  |  |  |  | LVEDD*, mm | LVEF†, % |
| Pt1 | 35 | M | 10‡ | 65 | 29 |
| Pt2 | 56 | M | 8§ | 56 | 40 |
| Pt3 | 52 | M | 9\|\| | 48 | 43 |
| Pt4 | 45 | M | 8# | 62 | 38 |
| Pt5 | 45 | F | 9** | 68 | 35 |
| Normal Values |  |  |  | <56 | ≥50 |

\* LVEDD=Left ventricular end-diastolic diameter
† LVEF= Left ventricular ejection fraction
‡= Emcitrabine/Tenofovir, Atazanavir, Ritonavir,
§=Emcitrabine/Tenofovir, Lopinavir/Ritonavir, Rivotril
||= Atazanavir, Norvir, Abacavir/Lamivudin

**Figure 1.**
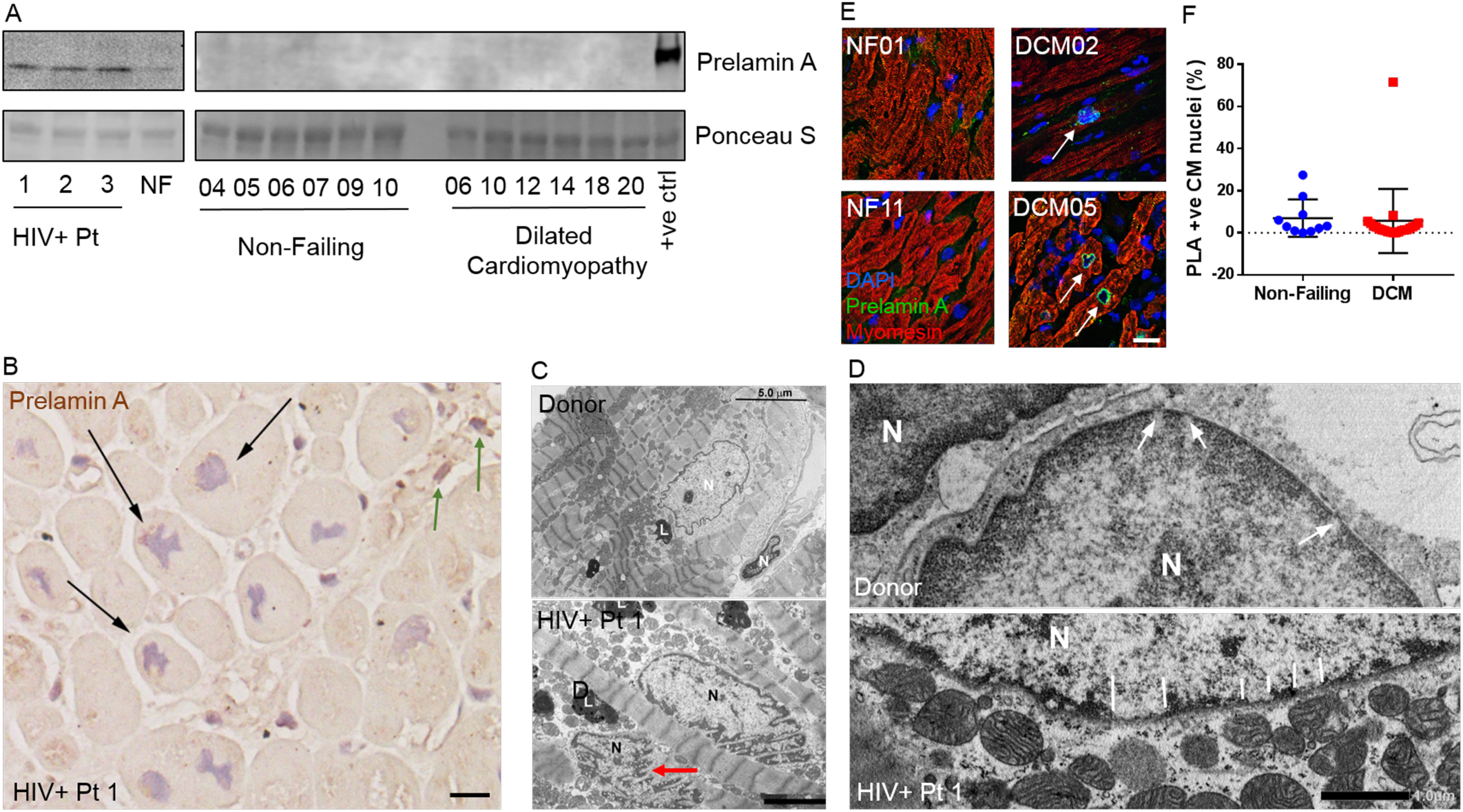
Prelamin A accumulates in hearts of patients with HIV-associated cardiomyopathy under retroviral therapy and Dilated Cardiomyopathy (DCM). **A.** Western blotting detected accumulation of prelamin A in hearts of HIV patients, but not in a selection of DCM patient samples. **B.** Immunohistochemistry showing focal prelamin A accumulation in CM nuclei (black arrows) and non-CM populations (green arrows) of HIV+ myocardium. Electron micrographs showing **C.** nuclear morphology defects in HIV+ myocardium (red arrow). Scale = 3 μm. **D.** Nuclear pore complexes surrounded by evenly spread heterochromatin in non-diseased myocardium (large white arrows) and heterochromatin displacement in HIV+ myocardium (small white arrows). Scale = 1 μm. N= Nucleus, L=Lipid bodies. **E**. Confocal micrographs of human heart sections from DCM patients and non-failing (NF) controls subjected to immunofluorescence staining to detect prelamin A (green), myomesin (red) and DAPI (blue). Arrows point to prelamin A positive CM nuclei, many of which exhibit nuclear morphology defects. **F.** the number of nuclei which stained positively for prelamin A were quantified as a percentage of CM nuclei for non-failing (blue circles, N=10) and DCM (red squares, N=21) myocardial sections (means ± SDs). Scale = 10 μm.

### Prelamin A accumulation in cardiomyocytes of mice causes cardiomyopathy and premature death by heart failure

Cardiomyocyte specific prelamin A transgenic (csPLA-Tg) mice (Fig. 2A) were born in a normal Mendelian ratio and were indistinguishable from floxed control (FLctrl) mice at birth. Immunofluorescence confirmed that accumulation of prelamin A in csPLA-Tg mice was specific to the nuclear rim of CMs (Fig. 2B) and was cardiac specific as shown by Western blot (Supp. Fig. 1).

**Figure 2.**
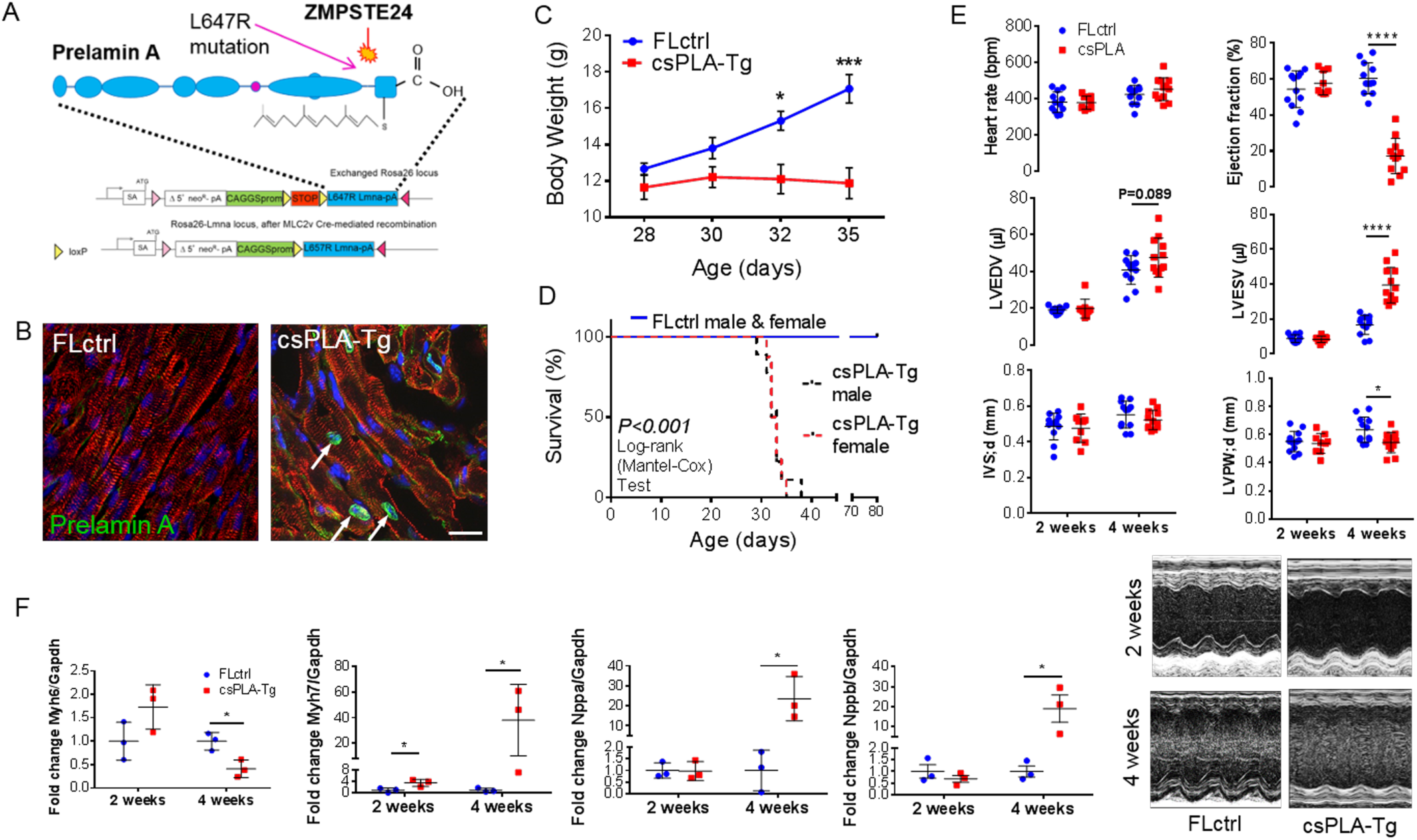
Targeted transgenesis of prelamin A led to nuclear ac cumulation in CMs and resulted in cardiomyopathy and premature death by heart failure in mice. **A.** schematic representation showing the site and of prelamin A (*LMNA*-L647R) cDNA insertion and the modifications required for conditional expression. **B.** Confocal micrographs of myocardium stained for prelamin A showing nuclear rim localisation in csPLA-Tg hearts. Scale = 10 μm. **C.** Growth curves showing that csPLA-Tg mice stop growing after 30 days (2-way ANOVA with Bonferroni multiple comparisons *P<0.05, ***P<0.001). **D.** Kaplan-Meier survival analysis showing that csPLA-Tg male and female mice die early compared to FLctrl counterparts (log-rank P<0.0001). **E.** Graphs of echocardiography analysis performed on movies acquired in B-mode showing severely compromised cardiac function in four-week old mice N=8-12/group. **F.** Foetal gene expression is dysregulated in csPLA-Tg hearts at four weeks indicating heart failure N=3/group. Values are means ± SDs. *P<0.05 **P<0.01 ****P<0.0001

After weaning (day 21) csPLA-Tg mice ceased to grow and died prematurely. By 32 days, body weight was significantly lower in csPLA-Tg mice (Fig. 2C) and median survival was significantly attenuated in male and female mice compared with FLctrl (Fig. 2D).

At two weeks csPLA-Tg mice showed no change in structural, dimensional or functional parameters by echocardiography as compared to FLctrl controls (Fig. 2E). In contrast, echocardiographic and MRI analysis of four-week mice showed that there was significant chamber dilatation as evidenced by increases in LV end systolic and diastolic volumes, and significant contractile impairment as evidenced by a marked reduction in ejection fraction. There was also evidence of LV posterior wall thinning. Heart rates were similar in the two groups (Fig. 2E, Supp. Fig. 2).

Quantitative (q)PCR analysis of csPLA-Tg myocardium showed that there was reduced mRNA expression of *Myh6* and increases in *Myh7*, *Nppa* and *Nppb* mRNA consistent with heart failure (Fig. 2F). This was supported by post-mortem analysis which showed csPLA-Tg hearts were enlarged at four weeks (Supp. Fig. 3A) whilst mass, based on heart weight relative to tibia length, was comparable between csPLA-Tg and FLctrl mice at two and four weeks (Supp. Fig. 3B). Transudative pleural effusions were evident upon opening the chest cavity in four-week old csPLA-Tg mice symptomatic of heart failure (Supp. Fig. 3C).

**Figure 3.**
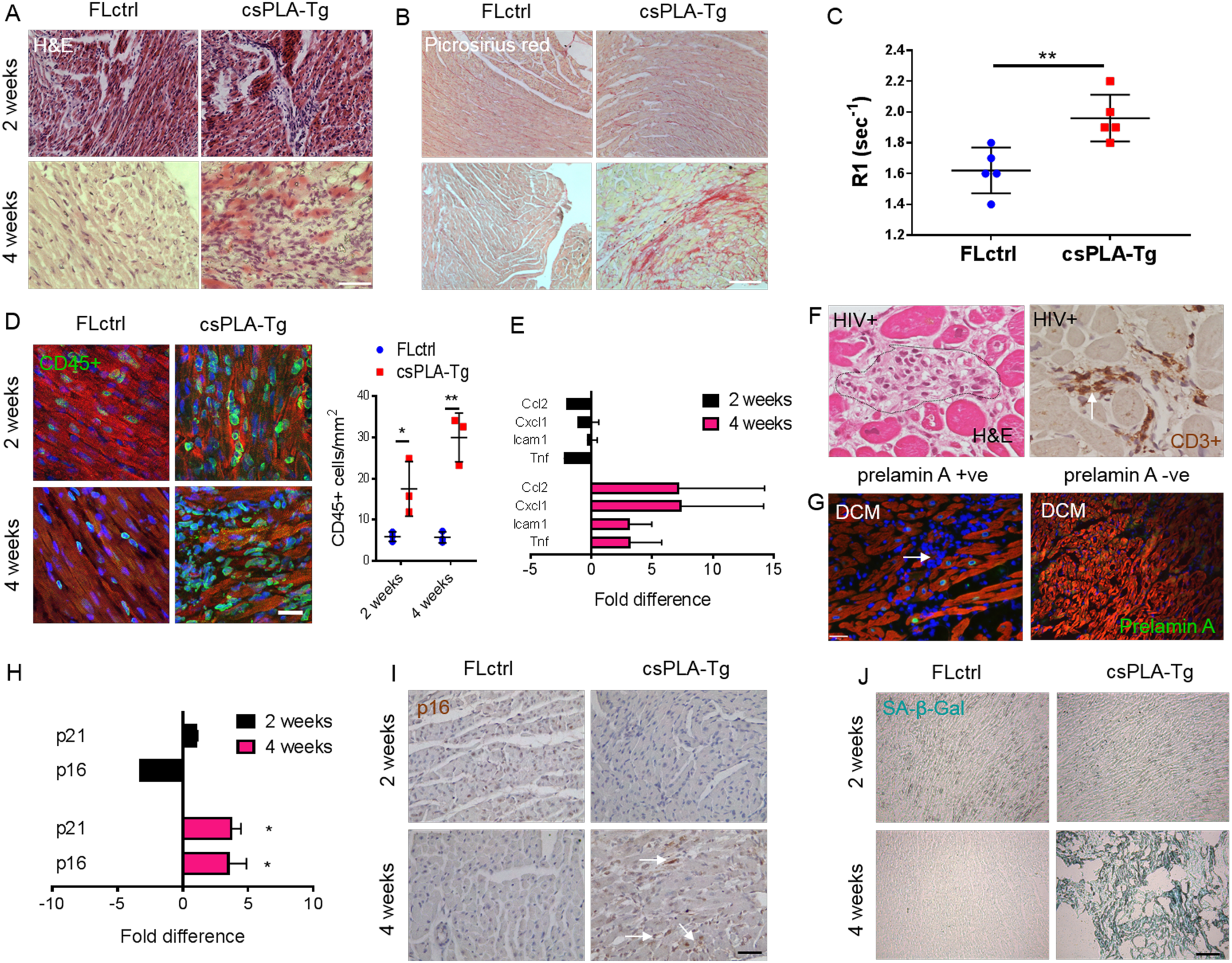
Fibrotic remodelling of csPLA-Tg myocardium occurred in tandem with inflammation and senescence whilst also sharing histological features with HIV associated cardiomyopathy and prelamin A mediated DCM. **A.** Light micrographs showing myocardial disarray in four week csPLA-Tg myocardium stained with Haematoxylin and Eosin. Scale = 30μm. **B.** Light micrographs showing Picrosirius red stained myocardium to indicate fibrosis in four week csPLA-Tg myocardium shown by excessive red staining. Scale = 30μm. **C.** Increased relaxation time (R1) of gadolinium contrast in four week csPLA-Tg myocardium indicative of fibrosis remodelling. **D.** Quantitative fluorescence immunostaining for CD45 shows presence of CD45+ cells in two and four-week csPLA-Tg myocardium. Scale = 10 μm. Values a re means ± SDs. N=3/group **P<0.01 **E.** qPCR showing the cytokine profile of csPLA-Tg myocardial mRNA. N=4/group **F.** H&E and CD3+ immunohistochemistry showing inflammation in HIV+ myocardium consistent with the csPLA-Tg model. Scale = 30μm. **G.** DCM patient sample in which Prelamin A accumulated showed mononuclear infiltration. Scale = 40 μm. **H.** expression of the mRNA for senescence markers p16 and p21 was elevated in four week csPLA-Tg hearts **I.** Immunohistochemical staining showing increase in p16 staining in 4 week csPLA-Tg myocardium. Scale = 30 μm. **J.** Senescence associated β galactosidase assay was performed to reveal bright blue staining in four week csPLA-Tg myocardium. Scale = 30 μm.

Enzyme linked immunosorbent assay (ELISA) of blood plasma identified a substantial increase in the plasma concentration of cardiac troponin T in four-week csPLA-Tg mice, indicative of significant CM damage or death (Supp. Fig. 4A). Increases in terminal deoxynucleotidyl transferase dUTP nick end labeling (TUNEL) positive nuclei indicated cell death occurred (Supp. Fig. 4B), though there was no evidence of caspase 3 cleavage or lamin cleavage, indicators for apoptosis (Supp. Fig. 4C), suggesting necrosis rather than apoptosis was the mode of cell death.

**Figure 4.**
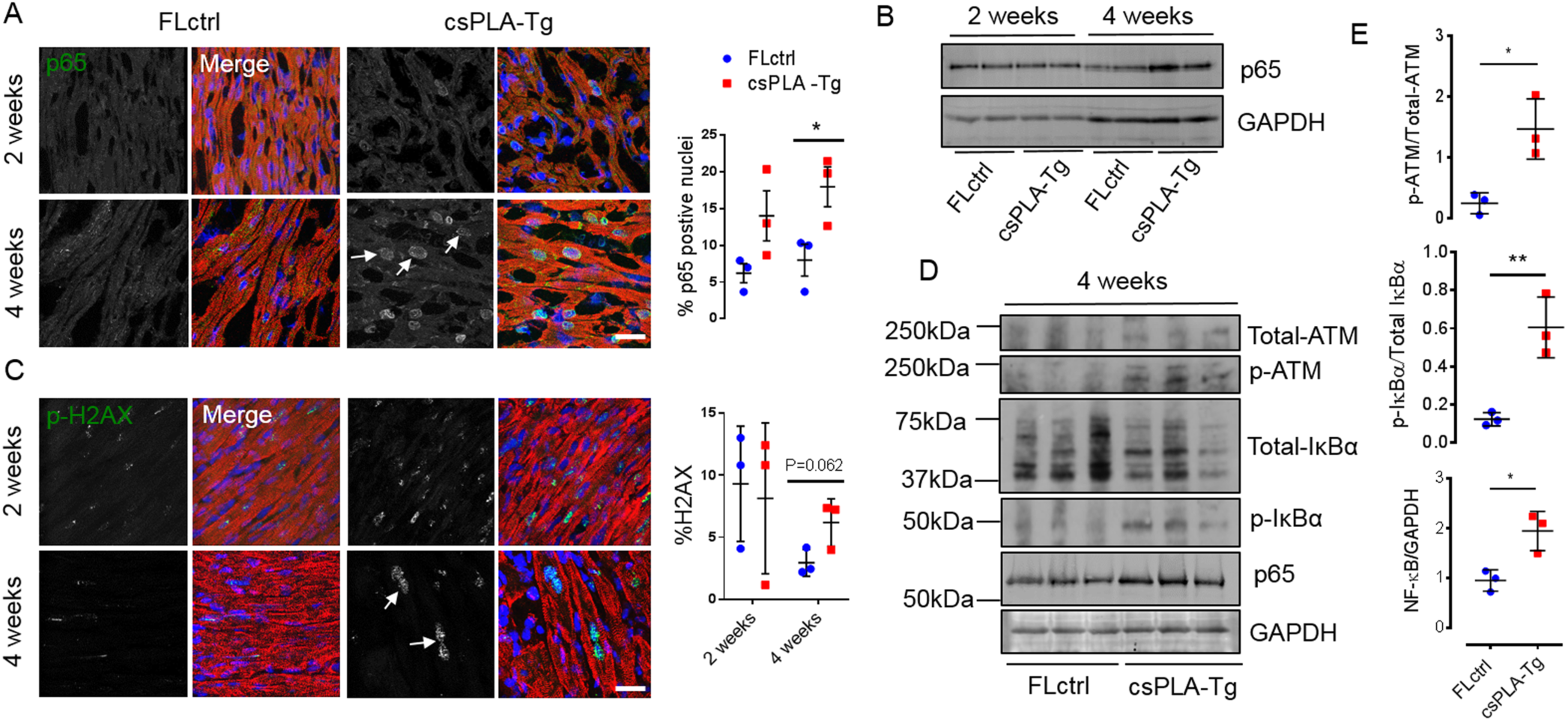
NF-κB signalling was activated in four week csPLA-Tg CMs and mediated by persistent DNA damage. **A.** Confocal micrographs of fluorescence immunostaining showing DNA damage marker γ-H2AX (white arrows) and quantification of γ-H2AX micrographs showing no significant changes between csPLA-Tg and FLctrl. **B.** Subcellular localisation of p65 subunit of NF-κB in csPLA-Tg myocardium, highlighted by white arrows and quantification of p65 micrographs showing increases in the number of nuclei expressing p65 for two and four week hearts counted as a percentage of total nuclei **C.** Western blot showing increase of NF-κB sub-unit p65 in four week csPLA-Tg myocardium. **D.** Western blots of four week myocardial lysates showing phosphorylation status of ATM and IκBα and **E.** graphs showing corresponding densitometry analyses. Scale = 10 μm. Values are means ± SDs. N=3/group *P<0.05 **P<0.001.

### csPLA-Tg myocardium exhibits fibrotic remodelling via a senescence associated secretory phenotype and mimics features of HIV associated cardiomyopathy

Consistent with the physiological data, haematoxylin and eosin (H&E) staining revealed that after perfusion fixation in diastole, csPLA-Tg heart chambers were normal at two weeks but dilated at four weeks (Supp. Fig. 3D, E). Inspection at higher magnification showed that csPLA-Tg heart tissue at two weeks was mostly normal, with sporadic regions of mononuclear aggregation in the myocardial interstitium. At four weeks however csPLA-Tg myocardium was in disarray and mononuclear infiltration was observed (Fig. 3A). Similarly, picrosirius red staining was comparable between csPLA-Tg and FLctrl at two weeks, but at four weeks csPLA-Tg myocardium showed substantial fibrosis as detected by picrosirius red staining and increased relaxation time of a gadolinium contrast agent in magnetic resonance imaging (MRI) of hearts (Fig. 3B,C). Observation of mononuclear infiltration in the myocardium suggested that inflammatory cells are activated and present in csPLA-Tg hearts and this was confirmed by immunofluorescence staining of myocardial sections for CD45 (Fig. 3A), which showed increased numbers of CD45+ cells in the myocardium of both two and four week old mice (Fig. 3B). mRNA expression analysis of myocardium for pro-inflammatory cytokines found that at four weeks *Tnf*, *Icam1*, *Cxcl1* and *Ccl2* were elevated (Fig. 3E). Myocardial inflammation has not previously been reported in models of *LMNA* cardiomyopathy, so to test whether this was unique to our model we performed immunostaining on *Lmna*^*-/-*^ myocardium for CD45 and observed no increase in CD45+ cells compared with wildtype suggesting this was a feature unique to prelamin A accumulation (Supp. Fig. 5). HIV+ cardiac tissue sections subjected to H&E staining and CD3+ immunohistochemistry confirmed inflammation within the myocardium (Figure 3E). We also observed regions of mononuclear infiltration in the DCM sample with perinuclear prelamin A which was not observed in other DCM samples (Fig. 3E). Moreover, electron microscopy of csPLA-Tg myocardium showed that heterochromatin displacement in CM nuclei could be observed as early as two weeks and nuclear morphology defects were present at 4 weeks (Fig. 3F).

**Figure 5.**
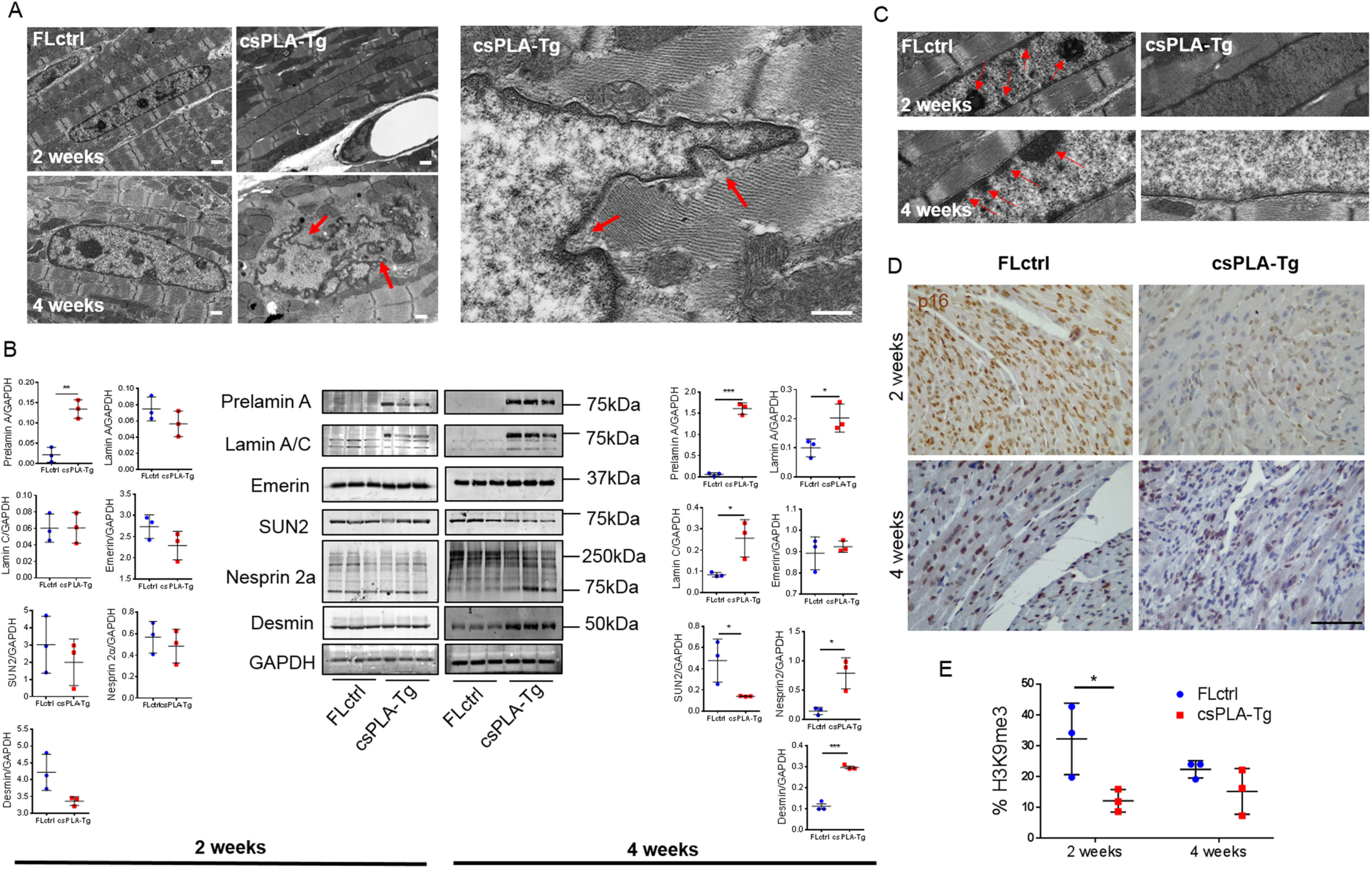
Prelamin A accumulation led to disorganisation of molecular structure at four weeks and loss of chromatin and histone marks at two weeks. **A.** Electron micrographs showing nuclear shape and size changes in csPLA-Tg myocardium, red arrows point to regions of nuclear in-folding characteristic of nuclear morphology defects. Scale = 500 nm. **B.** Western blot analysis showing the protein expression changes occurring at four weeks in structural proteins of the nuclear envelopelamin A/C, emerin, SUN2, nesprin 2, and also the cytoskeleton-desmin, with corresponding semi-quantitative densitometry analysis. Values are means ± SDs *P<0.05 **P<0.01 ***P<0.001. **C.** Electron micrographs show heterochromatin displacement and loss of chromocentres. Scale = 500 nm. D. Quantitative immunohistochemistry showed a profound loss of H3K9me3 staining as a percentage of total Haemotoxylin stain in two week csPLA myocardium. Scale = 30 μm. Values are mean ± SD. N=3/group. * P<0.05.

Because disruption to the nuclear lamina is associated with premature senescence (27, 28) and in turn senescence is associated with inflammation via the senescence-associated secretory phenotype (SASP) (29), we postulated that csPLA-Tg myocardium may display traits of senescence. Senescence-associated β-galactosidase assay showed that four-week csPLA-Tg myocardium displayed intense β-galactosidase expression when compared with FLctrl (Fig 3F). In addition, q-PCR showed expression of mRNA for the genes of senescence markers p16 *(Cdkn2a*) and p21 (*Cdkn1a*), were upregulated in four-week csPLA-Tg myocardium (Fig. 3E). Immunohistochemistry for p16 confirmed an increase at the protein level in four week csPLA-Tg myocardium (Fig. 6G).

**Figure 6.**
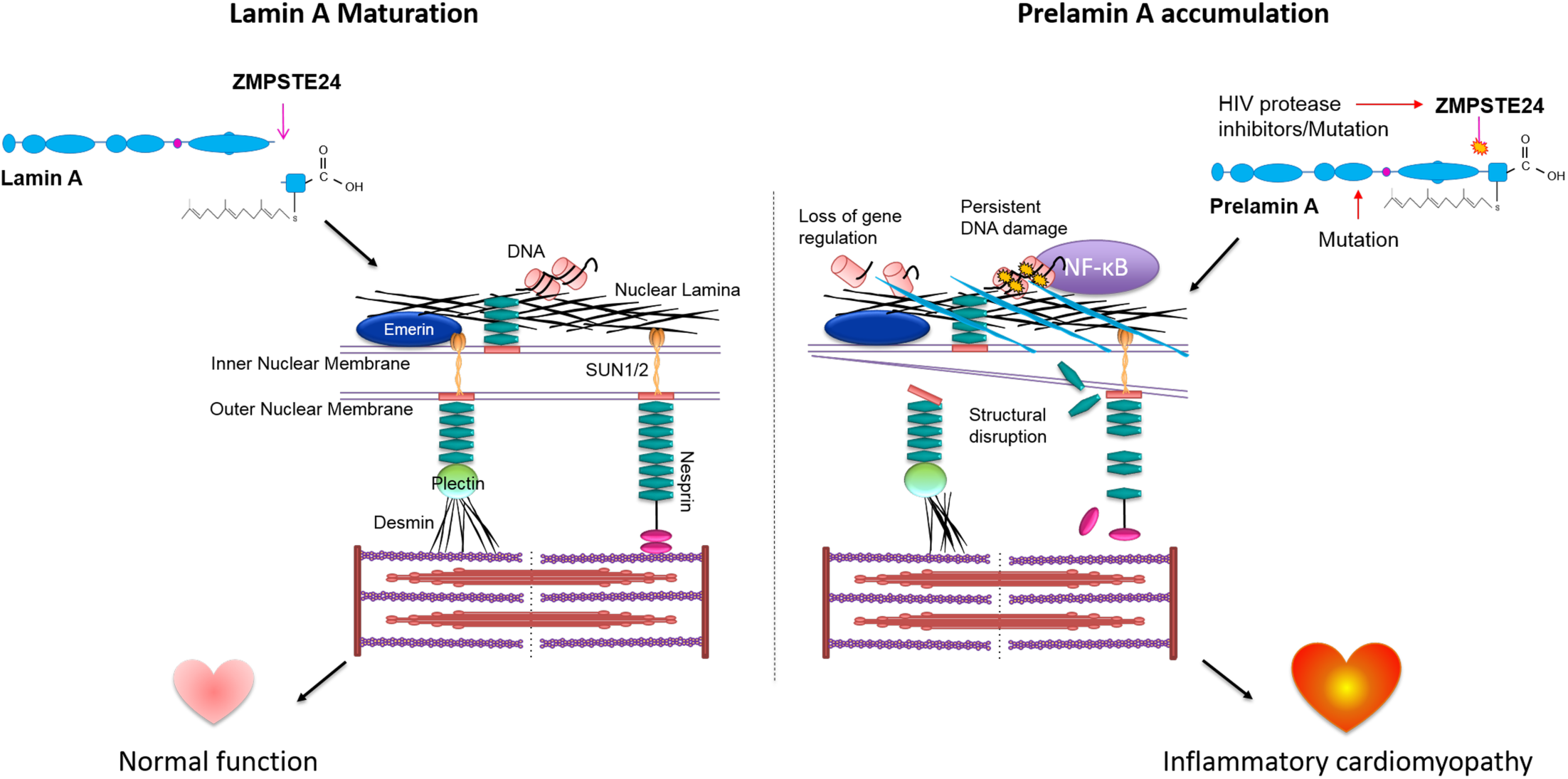
Schematic showing molecular and phenotypic implications of prelamin A accumulation in the heart. In a normally functioning heart prelamin A is cleaved by ZMPSTE24 leading to lamin A insertion in the nuclear lamina. Prelamin A accumulation may occur by mutations in *LMNA* and *ZMPSTE24* genes, and also by inhibition of ZMPSTE24 by HIV protease inhibitors. In this setting prelamin A accumulation is likely to contribute to the development of inflammatory cardiomyopathy. In the case of HIV PI inhibiton of ZMPSTE24, switiching treatments to HIV PIs with low- or no affinity to ZMPSTE24 may prove beneficial.

### Nuclear factor kappa-light-chain-enhancer of activated B cells (NF-κB) signalling was activated in csPLA-Tg myocardium

NF-κB is a master regulator of inflammation (30). It is also known that prelamin A can activate NF-κB via persistent DNA damage and non-canonical signalling pathway involving signalling partners such as IκBα (31). Persistent activation of the DNA damage response is also for activation of senescence in laminopathies (9). We hypothesised this might be activated in csPLA-Tg mice. The p65 subunit of NF-κB is translocated to the nucleus upon activation so to investigate this we performed quantitative immunofluorescence staining and showed that this was occurring at four weeks (Fig. 4A). To substantiate this finding Western blot indicated elevated expression of p65 at four weeks but not two weeks (Fig 4B). We then assessed DNA damage signalling by phosphorylated Histone 2AX (γ-H2AX)—a first responder and activator of DNA damage signalling. We found that γ-H2AX staining as a percentage of total nuclear stain was inconsistent in two week myocardium whilst in four-week csPLA-Tg myocardium there was a trend towards an increase (P = 0.062) (Fig. 4C). To investigate further we assessed the DNA damage transducer Ataxia Telangiectasia Mutated (ATM), which can activate NF-κB signalling via IκBα. Western blotting of myocardial lysates from four-week old mice showed ATM and IκBα were consistently phosphorylated in csPLA-Tg myocardium (Figs. 4D and E) Taken together these data infer that activation of inflammatory NF-κB signalling via ATM is a consequence of prelamin A accumulation in heart.

### Disruption to the LINC complex and cytoskeleton was preceded by loss of histone marks in csPLA-Tg myocardium

Inconsistencies between the activation of NF-κB and the onset of inflammation encouraged us to explore other pathways consistently affected in two week myocardium. The structural hypothesis of lamin dysfunction argues that nuclear envelope disruption can lead to increased susceptibility to mechanical stress and structural instability of cells (3). EM of csPLA-Tg myocardium showed, similar to HIV+ myocardium, that nuclear morphology defects could be observed at four weeks, though not at two weeks. Western blotting of LINC complex proteins and the intermediate filament desmin showed profound changes in expression at four weeks but not at two weeks (Fig 5B). Another theory of lamin dysfunction hypothesises that regulation of gene expression is affected by lamina disruption. EM images showed a loss of chromocentres and heterochromatin bundles which did appear to occur at two weeks in csPLA-Tg myocardium, again, supporting observations from HIV+ patient myocardium (Fig. 5C). We decided therefore to assess the chromatin changes by investigating methylation of lysine 9 of histone 3 (H3K9me3). We performed and quantified immunohistochemical staining for H3K9me3 expression and discovered a profound loss of H3K9me3 in two week csPLA-Tg myocardium (Fig. 5D, E).

## Discussion

### Accumulation of prelamin A occurs in genetic and HIV associated cardiomyopathy

To date the role of prelamin A in the setting of cardiomyopathy has been understudied. Here we present compelling evidence suggesting that prelamin A accumulation occurs in HIV associated cardiomyopathy, since all samples tested showed elevated prelamin A abundance, and nuclear morphology defects were observed. We also investigated a cohort of patient samples, for which the primary diagnosis was idiopathic DCM. We showed consistent prelamin A accumulation in one DCM sample and sporadic accumulation in all other samples. Although sequencing of this sample could not be performed, the consistent detection of prelamin A suggested there may be a genetic basis for the disease in this patient. Additionally, this accumulation is likely to exacerbate cardiomyopathy as evidenced by our study of a novel csPLA-Tg mouse line. A baseline phenotype of severe cardiac dysfunction was observed. The inflammatory response we observed, which correlates with the HIV associated cardiomyopathy phenotype, was also uniquely associated with myocardial senescence. Mechanistically, we linked the inflammatory response to modulation of NF-κB signalling, via activation of ATM. However, epigenetic changes may be more important when considering early initiating mechanisms.

### Prelamin A accumulation causes “Inflammageing” of the myocardium in mice

Though inflammation is known to occur during and after ischaemic events and later in heart failure (32) it is less commonly described in the progression to DCM. However, HIV associated cardiomyopathy is strongly associated with an inflammatory response in the myocardium (33, 34). Recent reports show that HAART exposure in perinatal cases of HIV ultimately benefits heart function compared with patients from the pre-HAART era (35) for whom opportunistic infections (36) and potential incorporation and replication of HIV into CMs were problematic (37). Nevertheless, a decline in cardiac function occurs when compared to the normal population and HAART therapies may contribute to this. NRTI’s, which prevent replication of HIV, also inhibit transcription of mitochondrial DNA and have been shown to cause cardiac dysfunction in mice (24, 25). Our study presents evidence that inflammation is the key outcome of prelamin A accumulation in CMs, and this is also a relatively unique phenotype of HIV associated cardiomyopathy. Though not subjected to rigorous interrogation of inflammatory pathways, global *Zmpste24*^*-/-*^ mouse myocardium also showed leukocyte infiltration further supporting a role for prelamin A toxicity in driving inflammation (38). In contrast, inflammation has not been reported in other *Lmna* mouse models of cardiomyopathy. Moreover, cardiomyocyte specific overexpression of wildtype lamin A in mice showed no phenotypic effect or impact on survival implying that processing mechanisms are able to cope with an increase in prelamin A concentration (39). In this context, it appears that accumulation prelamin A which cannot be processed, is pathogenic.

We provide evidence that the trigger for inflammation is likely linked to myocardial senescence. Whilst expression of γ-H2AX, the most commonly used marker for DNA damage, was inconsistent in csPLA-Tg myocardium, ataxia telangiectasia mutated (ATM) which signals down-stream of γ-H2AX, was persistently phosphorylated at four weeks. We observed that p16 and p21 mRNA was increased, along with mRNA of pro-inflammatory cytokines Tnf-α, Icam1, Cxcl1, Ccl2, suggesting that myocardium in these mice exhibit the SASP. We were able to further substantiate this by the detection of senescence associated β galactosidase (SA-β-Gal) in csPLA-Tg myocardium. Furthermore, these data concur with previous work showing that when prelamin A accumulates in vascular smooth muscle cells activation of the SASP occurs (40).

### Early loss of repressive histone marks indicates gene expression pathways are initiators of pathogenesis

Global *Zmpste24*^*-/-*^ mice suffer from systemic inflammation arising from non-canonical ATM-dependent NF-κB signalling (31) and we showed that this pathway was activated locally in csPLA-Tg hearts at four-weeks making it likely that increases in SASP factors are caused by activation of NF-κB. However, the absence of NF-κB signalling at two weeks suggests that this inflammatory pathway propagates rather than initiates disease mechanisms. Therefore, we speculated to other mechanisms for disease genesis. One of these was based on the mechanical hypothesis of lamin dysfunction. We reasoned that susceptibility to mechanical stress, which is continual and repetitive in the heart, might lead to structural defects in the nuclear envelope and cytoskeleton. However, these were not observed until four weeks, meaning this was unlikely. Therefore we speculate to the existence of a priming mechanism for the initiation of disease remodelling related to epigenetic changes, since chromatin displacement and loss of H3K9me3 were observed in two week mouse hearts. The mechanisms downstream of this remain to be determined but may involve misregulation of the polycomb repressive complexes (41), known to control senescent genes such as those encoding p16 and p21 and potentially induce SASP independently of DNA damage pathways (42).

## Conclusions

In summary we have identified a novel role for prelamin A in HIV associated and dilated cardiomyopathies. Accumulation of prelamin A has catastrophic consequences for the integrity of the myocardium resulting in an “inflammageing” phenotype and subsequent loss of contractility. Whilst targeting inflammation potentially via ATM activity may prove a useful for patients with established DCM owing to prelamin A accumulation, the immediate translational aspect of this study lies in the implications for the treatment of HIV associated cardiomyopathy patients, for whom, a change of therapy may have a beneficial outcome in the clinic. Elegant biochemistry performed by Robinson and colleagues provided compelling evidence that HIV PIs bind and block activity of ZMPSTE24 (43). Moreover they were also able to show a rank order of affinity to ZMPSTE24 of HIV PIs currently available for use-lopinavir > ritonavir > amprenavir > darunavir. Of these, darunavir was shown not to bind *ZMPSTE24* at all, which confirms earlier work (44). Many HIV+ patients suffering from cardiac symptoms are not currently using darunavir in their HAART regimes (Table 1). Adjusting HAART regimes to incorporate darunavir and other HIV protease inhibitors with low affinity to ZPMSTE24 may reduce prelamin A accumulation and provide therapeutic benefit for patients suffering from symptoms of HIV associated cardiomyopathy.

## Materials and Methods

### Human studies

This study complies with the declaration of Helsinki. Human DCM specimens were obtained from the Sydney Heart Bank (Hospital Research Ethical Committee approval #H03/118; University of Sydney ethical approval #12146) and from Papworth tissue bank in Cambridge, UK and used in accordance with ethical guidelines of King’s College London (REC reference 13/LO/1950) and the current UK law. Studies involving HIV associated cardiomyopathy patients endomyocardial biopsies were approved by the Ethics Committee of La Sapienza University Rome. All patients gave informed consent.

### Human Echocardiography

Echocardiographic parameters were determined according to established criteria. In particular, EF (ejection fraction) was calculated in the apical 4 and 2-chamber views from three separate cardiac cycles using the modified Simpson’s method and LVEDD (left ventricular end-diastolic diameter) was measured in long-axis and short-axis view.

### Generation of cardiomyocyte (CM) specific prelamin A transgenic (csPLA-Tg) mice

All animal procedures were performed in accordance with the Guidance on the Operation of the Animals (Scientific Procedures) Act, 1986 (UK Home Office). This mouse model was commissioned from Taconic-Artemis with the aim to devise a transgenic system to assess the *in vivo* effects of uncleavable prelamin A overexpression. We performed site directed mutagenesis of the human *LMNA* gene at leucine 647 and replaced it with arginine (*LMNA*-L647R). This corresponds to the cleavage site for *ZMPSTE24* and blocks cleavage. The system involved recombinase mediated cassette exchange (RMCE) of the Rosa26 gene whereby *LMNA-*L647R cDNA was inserted into an exchange vector containing a neomycin resistance gene, a strong CAGGS promoter sequence and a STOP cassette flanked by loxP sites. Electroporation into the embryonic stem (ES) cells of C57BL/6 mice led to site-specific recombination by the recombinases F3 and FRT. Neomycin resistant clones that had undergone RMCE were selected. After administration of hormones, superovulated BALB/c females were mated with BALB/c males. Blastocysts were isolated from the uterus at 3.5 days post coitum (dpc) for microinjection. Blastocysts were placed in a drop of DMEM with 15% FCS under mineral oil. A flat tip, piezo actuated microinjection-pipette with an internal diameter of 12-15 µm was used to inject 10-15 targeted C57BL/6NTac ES cells into each blastocyst. After recovery, 8 injected blastocysts were transferred to each uterine horn of 2.5 dpc, pseudopregnant NMRI females. Chimerism was measured in chimeras (G0) by coat colour contribution of ES cells to the BALB/c host (black/white). Highly chimeric mice were bred to strain C57BL/6 females. Recombination by mating with mice carrying cre-recombinase under the control of the myosin light chain 2 ventricular (MLC2v) promoter led to removal of the stop cassette allowing expression of uncleavable prelamin A in nuclei of CMs of affected offspring. These mice were called CM specific prelamin A transgenic (csPLA-Tg) mice. They were compared to mice expressing the transgene but retaining the lox P sites, termed floxed controls (FLctrl). All transgenic mice used in the study were heterozygous for the transgene and generated on a C57Bl6 background. Male and female mice were used in this study.

### Murine echocardiography

Echocardiography was performed using a Vevo® 2100 imaging system with a 30 MHz linear transducer specially designed for small animal studies (VisualSonics, USA). Echocardiography was performed with 5% isofluorane fast induction of anaesthesia followed by maintenance of 1-1.5% isoflurane anaesthesia for 4 week old mice and 2.5% isofluorane for 2 week old mice, which was vaporized in 100% oxygen delivered at 1.5-2 liters/min. Heart rate was kept at ∼400-450 beats per minute while respiratory rate was ∼100 breaths per minute. Body temperature was ∼36.5±1°C.

### Murine cardiac Magnetic Resonance Imaging (MRI) for T1 mapping

For detection of myocardial ‘scarring’ by T1 mapping (45), anaesthetised mice were subject to MR imaging before and 25 min after intraperitoneal (i.p.) administration of 0.75 mmol/kg of gadofosveset trisodium (Ablavar®, Lantheus Medical Imaging, North Billerica, MA), a gadolinium-based contrast agent. An ECG triggered, single slice, Look-Locker acquisition was used for T1 mapping and to measure R1 values of the myocardium. The slice was selected in the middle of the heart. Imaging parameters included FOV = 25 x 25 mm^2^, slice thickness = 1 mm, matrix size = 128 x 128, 3 phases/cycle, total of 30 phases, 1 slice, flip angle = 10°, TR. 2700 ms, TReff ≈ 40 ms ((cardiac cycle)/(3 phase/cycle)), TE = 2 ms, cardiac cycle = 120 ± 20 ms, number of averages = 1, acquisition time ≈ 13 min. T1-weighted sequences were analysed to assess the R1 values of the myocardium.

Look-Locker T1 mapping resulted in 30 images (3 per cardiac cycle) from which R1 values of blood, infarcted, remote and healthy myocardium were calculated using an exponential 3 parameter fit (A-B*exp (-TI/T1*)) with subsequent T1 correction (OriginLab Corporation, Wellesley, USA).

### Transmission Electron Microscopy (TEM)

Mice were injected intraperitoneally with heparin (5000 u/kg body weight). This was followed by intraperitoneal injection of 50mg/kg body weight of sodium pentobarbital to induce terminal anaesthesia. The chest cavity was opened and secured with a hemostat. The LV was injected with a needle connected via a pump to a reservoir of pre-wash buffer. Flow rate of the pump was adjusted as to perfuse mouse heart at a pressure between 90-100 mmHg with pre-wash buffer. Pre-wash buffer was a standard physiological tyrode solution containing 10mM BDM to arrest the heart in diastole and 2.5% PVP to replace the protein content of blood, thereby maintaining colloidal pressure and preventing haemorrhage at vascular sites in the heart. Following prewash, the hearts were perfused with fixative solution containing 2% Glutaraldehyde and 2% Paraformaldehyde until 20ml of fixative had been perfused. Hearts were dissected and the mid LV was isolated and cut for further processing. Samples were dehydrated through a graded series of ethanol washes and embedded in epoxy resin. Semi-thin sections (0.2 μm) were stained with toluidine blue for light microscopy examinations and were used to guide sampling for TEM studies. Thin sections (0.09 μm) were collected on 150-mesh copper grids and double stained with uranyl acetate and lead citrate for examination under TEM (H7650, Hitachi, Tokyo, Japan).

Human HIV+ samples were fixed in 2% glutaraldehyde in a 0.1 M phosphate buffer, at pH 7.3, post fixed in osmium tetroxide and processed following a standard schedule for embedding in Epon resin. Semi-thin sections were stained either with azur-II or basic fuchsin solutions and mounted with permount medium. Ultrathin sections (70-80) were stained with uranyl acetate and lead hydroxyde. A Jeol 1400 plus TEM was used for observation and photographic analysis.

### Statistics

All *in vivo* and *ex vivo* data of age-matched csPLA-Tg versus FLctrl mice were analysed using the Student’s unpaired T-test for normal distribution. Where standard deviations were substantially different Welsch’s correction was applied and in the case of non-normal distribution of data the non-parametric Mann-Whitney test was applied. For the analyses of body weights at over time, two-way analysis of variance with repeated measures was selected with Bonferroni post hoc test for multiple comparisons. Kaplan-Meier survival curves were assessed by the log-rank Cox-Mantel test. Values were expressed as means ± the standard deviation (SD). Tests were performed in Excel (Microsoft) or Prism (GraphPad).

Detailed accounts of other methods used in this study can be found in the online supplement to this article.

## Supporting information

## Acknowledgments

We would like to acknowledge the Papworth tissue bank (Cambridge, UK) for providing explanted DCM samples.

