## Supplementary material for "Accumulation of prelamin A drives inflammation in the heart with implications for treatment of inherited and acquired cardiomyopathies"

**Supplemental Materials for**

### 'Accumulation of prelamin A in cardiomyocytes drives inflammation in the heart with implications for treatment of HIV associated cardiomyopathy'

**Authors:** Daniel Brayson, PhD[1]; Andrea Frustaci, MD[2,3]; Romina Verardo, PhD[3]; Cristina Chimenti, MD PhD[2,3]; Matteo Antonio Russo, MD[4]; Robert Hayward, BSc[1]; Sadia Munir Ahmad, MSc[1]; Gema Vizcay-Barrena, PhD[5], Andrea Protti, PhD[6]; Peter S. Zammit, PhD[7], Cristobal G. dos Remedios, PhD[8]; Elisabeth Ehler, PhD[1,7]; Ajay M. Shah, MD[1]; Catherine M. Shanahan, PhD[1]

**Addresses:**

[1]School of Cardiovascular Medicine and Sciences, King's College London BHF Centre for Research Excellence, London, UK

[2]Department of Cardiovascular, Nefrologic, Anestesiologic and Geriatric Sciences, La Sapienza University of Rome, Italy.

[3]National Institute for Infectious Diseases IRCCS 'L. Spallanzani', Rome, Italy

[4]MEBIC Open University San Raffaele and IRCCS San Raffaele Pisana, Laboratory of Molecular and Cellular Pathology, Milan, Italy

[5]Centre for Ultrastructural Imaging, Guy's Campus, King's College London, London, UK

[6] Department of Imaging, Lurie Family Imaging Center, Center for Biomedical Imaging in Oncology, Dana-Farber Cancer Institute, Harvard Medical School, Boston, Massachusetts, USA,

[7]Randall Centre for Cell and Molecular Biophysics, King's College London, UK

[8]Department of Anatomy, Bosch Institute, University of Sydney, Sydney, NSW, Australia

**Working Title:** Brayson et al., Accumulation of prelamin A in cardiomyopathy

**Address for correspondence:** 3rd Floor James Black Centre
125 Coldharbour Lane
King's College London
London SE5 9NU

**Email:** daniel.brayson @kcl.ac.uk

**Supplemental Methods**

**Cardiac Magnetic Resonance for functional parameters**

Cardiac MRI was performed on a 7 Tesla (T) horizontal MR scanner (Varian Inc., Palo Alto, CA) with mice in the prone position. The gradient coil had an inner diameter of 12 cm; the gradient strength was 1000 mT/m (100 G/cm), and rise-time was 120 ms. A quadrature transmit/receive coil (RAPID Biomedical GmbH, Germany) with an internal diameter of 39 mm was used. Anesthesia was maintained with 1.5% isoflurane/98.5% oxygen and body temperature was maintained at 37°C using a warm air fan (SA Instruments, Stony Brook, NY). The ECG was monitored by means of two metallic needles placed subcutaneously in the front paws. A pressure-transducer for respiratory gating was placed on the animal abdomen. To synchronize data acquisition with the ECG and to compensate for respiratory motion, simultaneous ECG triggering and respiration gating (SA Instruments) were applied. Functional and volumetric parameters were achieved following a multi slice Cine-FLASH acquisition. These parameters were used: FOV = 25 x 25 $mm^2$, slide thickness = 1 mm, matrix size = 128 x 128, 9 to 10 frames/cycle, 9 slices, flip angle =40°, cardiac cycle = 120 ± 30 ms, averages = 3, acquisition time ≈ 8 min. Functional and volumetric parameters were calculated from cine-FLASH images and areas of contrast-enhancement were calculated using a semi-automated in-house developed computer software program (King's College London, ClinicalVolumes).

**Morphometry**

Mice were subjected to observation and weighed regularly. When it became apparent that the phenotype of the csPLA-Tg mice was failure to thrive and premature death, it was decided that when the weight of a mouse fell below 85% of its FLctrl littermates, it was deemed to be suffering the maximum threshold of pain permitted on the project license and was culled. Post-mortem morphometry analysis was performed using heart weight and tibia length.

**Indirect immunofluorescence of cryopreserved tissue**

Tissue was removed from storage at -80°C and warmed to -20°C for approximately 1 hour. Tissue was then mounted onto the stage of a cryostat with OCT compound and allowed to equilibrate to the ambient temperature of the cryostat for 5 min; for cutting heart tissue, the stage temperature was set to -22°C and the knife was set to -20°C. The sections were then cut to 10 µm thick and mounted onto high quality polylysine or superfrost slides. The sections were allowed to air dry for approximately 1 hour and then stored at -80°C. For staining, sections were allowed to dry completely and fixed by immersion, in a Coplin jar, in pre-chilled 100% methanol at -20°C for 5 min or 4% formalin for 10 min RT. Sections were washed 2x5 min in PBS. Sections were permeabilised in 0.5% NP-40 for 3 min RT and washed 3x5 min in PBS. Sections were blocked in 3% serum originating from a species to which the antibody was not raised against for 1 hour RT. Primary antibody was diluted appropriately in blocking solution applied to the sections and incubated in a humidified chamber overnight at 4°C. Sections were washed 3x5 min in PBS and secondary antibody conjugated to a fluorophore was diluted as appropriate, applied to the sections and incubated in the dark in a humidified chamber for 1 hour RT. DAPI was added 1:10,000 for 5 min at the end of the incubation for visualisation of nucleic structures. Sections were washed 3x5 min in PBS in the dark and then mounted in mowiol mounting media and allowed to dry in the dark overnight. Antibody dilutions can be found in Supplemental Table 1.

**Microscopy**

Immunohistochemical and general tissue staining was analysed by bright-field light microscopy using an Axioskop 2 microscope (Carl Zeiss MicroImaging, NY). Immunofluorescence was analysed using wide-field fluorescence microscopy (IX81 microscope, Olympus). Where images captured were subjected to quantitative analysis, this was performed by counting by eye or by selction and measuring tools made freely available by ImageJ software. In each case the investigator performing the analysis was blinded to group assignment. Images for publication were captured using a Leica SP5 confocal microscope (Leica Microsystems, UK). Confocal images were processed in Photoshop CS3

(Adobe) and minimal adjustments to brightness and contrast were made where necessary, e.g. when images were deemed too dark for comfortable viewing.

**Immunoblotting**

Cryopreserved heart tissue was homogenised by placement into a pre-liquid nitrogen cooled stainless steel chamber followed by crushing administered by a pre-liquid nitrogen cooled stainless steel rod and rubber mallet. 200 µl lysis/sample buffer (3.7 M urea, 134.6 mM Tris pH 6.8; 5.4% SDS; 2.3% NP-40; 4.45% beta-mercaptoethanol; 4% glycerol and 6 mg/100 ml bromophenolblue) was added. And frozen pellets placed into Eppendorf tubes. The tubes were heated to 85°C for 5 minutes. A chamber was prepared with an SDS gel of 10% acrylamide concentration. 20µl was then loaded into each well of the gel. The proteins were separated at 150 mV. Transfer was achieved in wet conditions. The proteins were subject to electrophoretic transfer onto PVDF membranes that had been incubated in methanol for 20 seconds and then washed in distilled water. The transfer conditions were 65 Amps at 4°C ON. The membrane was blocked for 1 hour with 5% Milk in TBS-T. Primary antibodies were added (Table 2.2) with TBS-T/5% milk or TBS-T 5% bovine serum albumin and incubated overnight at 4°C. The membrane was washed 3x5 min with TBS-T. The blots were then incubated in secondary antibody coupled to horseradish peroxidise for 1 hour at room temperature or secondary antibody conjugated to a fluorophore for 1 hour. The blots were again washed with TBS-T and bound antibody signal was detected by an Odyssey® scanning detection system (LI-COR Biosciences, U.S.A). Western blots were quantified using ImageStudio™ software (LI-COR Biosciences, U.S.A).

Immunoblotting of HIV associated cardiomyopathy tissue was performed using commercially available kit for whole-cell extraction from frozen tissue according to manufacturers instructions (Active Motif, USA, cat. no. 40010).

**Quantitative (q)-PCR**

2 µg RNA was diluted into 10 µl of double distilled (dd) $H_2O$. 0.3 µl Oligo dT primer, 0.3 µl random primer and 1µl dNTPs was added to this and heated to 65°C and then cooled to 4°C in a thermocycler. 4 µl 5x Buffer, 3.9 µl DEPC-treated $H_2O$ and 0.5 µl reverse transcriptase (M-MLV) was added and

samples were heated in a thermocycler to 25°C for 10 min, 37°C for 50 min and cooled to 4°C. After the reaction, the samples were diluted to 100 µl total volume for qPCR. qPCR was carried out using the ΔCt and standard curve methods. When the ΔCt method was used the primers had been pre-validated and were part of a standard protocol for the lab. 9 µl cDNA was added to 10 µl 2x Sybr green PCR master mix and 5 µmol of forward and reverse primers (Supp table. 2). Cycling parameters were 94°C for 15 seconds, followed by single-step annealing and extension at 60°C for 1 min (35 cycles). Reactions were carried out in a Corbett RotorGene-3000. The cycle threshold (Ct) was determined manually as an arbitrary point during the linear phase of amplification and mRNA expression was quantified as a ratio of GAPDH expression. mRNA expression of csPLA-Tg groups was expressed as fold change compared to FLctrl groups.

**Haemotoxylin and eosin staining**

Tissue samples were fixed in 4% PFA and dehydrated overnight. The samples were then embedded in wax blocks and cut to sections of 6 µM thickness and mounted on glass slides (superfrost plus, VWR International). Slides were baked at 60°C overnight. Tissues were then processed and rehydrated by sequential incubation in xylene and reducing concentrations of ethanol 100%, (2x3 min) 96%, 75%, 50%, 25% (1 min). After rehydration, slides were left under running tap water for 10 min. After rehydration, sections were washed under running tap water for 5 min, and then placed in Harris' haematoxylin solution for 5 min. The sections were then washed under running tap water until the water ran clear followed by differentiation in acid alcohol. The sections were again placed under running tap water for 5 min and then stained with eosin solution for 3 min. The sections were dunked 5 times in tap water and then subjected to dehydration by sequential ethanol treatment: 25%, 50%, 75%, 96% (1 min each) 100%, (2x3 min) xylene, (2x5 min). Coverslips were mounted using DPX and allowed to dry overnight.

**Picro-sirius red staining**

Following rehydration, slides were left in $ddH_2O$ for 5 min followed by a 30 second incubation in 0.2% phosphomolybdic acid. Slides were rinsed in $ddH_2O$ and then left in 1% Sirius red solution for

90 min. Slides were washed 2x2 min in acidified dd$H_2O$ (0.05% acetic acid) followed by a 15 min incubation in Picric acid. Slides were rinsed several times in dd$H_2O$ followed by dehydration by sequential ethanol treatment: Glass coverslips were mounted with DPX and allowed to dry overnight.

**Senescence associated β-Galactosidase assay**

Hearts were resected from mice and immediately frozen and cryosectioned into 5mm thick slices and mounted onto superfrost slides. Sections were allowed to dry for 1-2 hrs. Samples were fixed in 1% formalin solution for 10 min RT. Senescence associated β-Galactosidase (SA-β-Gal) staining solution was freshly made and applied to each heart section so that they were generously submerged (approx. 50 µl per heart section). The slides were incubated in a humidified chamber overnight (approx. 16h) at 37°C. The slides were washed 2x5 min in PBS and once in 100% methanol for 2 min, the heart sections were then viewed and photographed under bright-field microscopy.

**Cardiac troponin T (cTnT) Enzyme Linked Immuno-Sorbent Assay (ELISA)**

Cardiac puncture under terminal anaesthesia was performed and blood was collected in an EDTA coated Eppendorf and spun at 13,000 g 4°C for 15 minutes to sediment red blood cells. Plasma was collected and subject to ELISA for cTnT (Elabscience, USA) according to manufacturer's instructions.

**TUNEL staining**

Deparaffinised sections were digested in 20 µg/ml proteinase K solution at 37°C for 10 min and washed 3x5 min in PBS. Sections were then incubated in 3% $H_2O_2$ for 10 min RT to block endogenous peroxidase activity and washed 3x5 min in PBS. Sections were then pre-incubated in TdT reaction buffer for 10 min RT and then incubated in TdT reaction mixture for 1 hour at 37°C in a humidified chamber. Sections were then washed in stop wash buffer for 10 min RT and rinsed in PBS-T 3x5 min. Sections were incubated with streptavidin-HRP for 20 min RT. Sections were incubated in DAB chromagen solution for 1-10 min (until brown staining appeared) and left under running tap water for 5 min. Coverslips were mounted with DPX and allowed to dry for 2 hour or overnight (ON).

**Supplemental Tables**

**Supp Table 1. List of antibodies used in this investigation**

| **Primary Antibodies** | **Source** | **Dilution (WB)** | **Dilution (IF)** | **Dilution (IHC)** |
|---|---|---|---|---|
| CD3 | Dako M7254 | - | - | 1:20 |
| CD45 | Abcam ab10558 | 1:1000 | 1:100 | - |
| Cleaved Caspase3 | Cell Signaling 9661 | 1:1000 | - | - |
| Desmin | Dako M0760 | 1:100 | - | - |
| Emerin | Leica Biosystems NCL-Emerin | 1:500 | - | - |
| GAPDH | Sigma-Aldrich G8795 | 1:1000 | - | - |
| H3k9me3 | abcam | | | 1:1000 |
| Lamin A/C (N-18) | Santa Cruz Biotechnology sc-6215 | 1:1000 | - | - |
| Myomesin B4 | C/O EE (homemade) | - | 1:100-200 | - |
| Nesprin 2 N3 | Homemade | 1:1000 | - | |
| γ-H2AX | Cell signalling 9718 | 1:1000 | 1:1000 | - |
| p16 | Cell signalling 4824 | | - | 1:1000 |
| p65 | Santa cruz Biotechnology sc-372 | 1:500 | 1:200 | - |
| Prelamin-A (C-20) | Santa Cruz Biotechnology sc-6214 | 1:200 | 1:1000 | |
| SUN 2 | Abcam ab65447 | 1:100 | - | - |
| **Secondary Antibodies** | **Source and cat number** | **Dilution (WB)** | **Dilution (IF)** | **Dilution (IHC)** |
| Anti-goat Alexa fluor 488. | Invitrogen A-11055 | - | 1:1000 | - |
| Anti-goat Alexa fluor 568 | Invitrogen A-11057 | - | 1:500 | - |
| Anti-mouse Alexa 488 | Invitrogen A11004 | - | 1:1000 | - |

| | | | | |
|---|---|---|---|---|
| Anti-mouse Cy3 | Jackson Laboratories 715-150-150 | - | 1:1000 | - |
| Anti-rabbit Alexa fluor 488 | Invitrogen A-11011 | - | 1:1000 | - |
| Biotinylated anti-goat | VECTASTAIN ABC kit (see reagents) | - | - | 1:500 |
| IRdye 800 Anti-goat | LICOR 926-32214 | 1:15000 | - | - |
| IRdye 800 Anti-mouse | LICOR 926-32210 | 1:15000 | - | - |
| IRdye 800 Anti-rabbit | LICOR 926-32211 | 1:15000 | - | - |
| IRdye680 Anti-rabbit | LICOR 926-68073 | 1:15000 | | |

**Supp Table 2. Primers used in this investigation for qPCR experiments**

| Primers | Sequence (5’-3’) |
|---|---|
| Nppa | F—CGTGCCCCGACCCACGCCAGCATGGGCTCC<br>R—GGCTCCGAGGGCCAGCGAGCAGAGCCCTCA |
| Nppb | F—AAGGGAGAATACGGCATCATT<br>R—ACAGCACCTTCAGGAGATCCA |
| Myh6 | F—CAGAGATTTCTCCAACCCAGCTGCG<br>R—AGTCAGCCATCTGGGCGTCCG |
| Myh7 | F—AGCAGCAGTTGGATGAGCGACT<br>R—CCAGCTCCTCGATGCGTGCC |
| GAPDH | F—CGTGCCGCCTGGAGAA<br>R—CCCTCAGATGCCTGCTTCAC |
| Ccl2 | F—TTAAAAACCTGGATCGGAACCAA<br>R—GCATTAGCTTCAGATTTACGGGT |
| Icam1 | F—GGCATTGTTCTCTAATGTCTCCG<br>R—TGTCGAGCTTTGGGATGGTAG |
| Tnf | F—CAGGCGGTGCCTATGTCTC<br>R—CGATCACCCCGAAGTTCAGTAG |
| Cxcl1 | F—ACTGCACCCAAACCGAAGTC<br>R—TGGGGACACCTTTTAGCATCTT |
| Cdkn2a (p16) | Qiagen (Cat |
| Cdk1a (p21) | Qiagen (Cat |

**Supplemental Table 3.** Patient information for explanted DCM samples

| Random ID | Assigned ID | Sex | Age | Clinical Diagnosis | Gene | Mutation | Ethnicity | Diabetes | Ejection fraction (%) |
|---|---|---|---|---|---|---|---|---|---|
| 2.059 | DCM01 | M | 43 | FDCM | NA | NA | NA | NA | NA |
| 4.032 | DCM02 | M | 31 | DCM | NA | NA | NA | NA | NA |
| 4.047 | DCM03 | F | 63 | FDCM | MYOM1 | E247K | NA | NA | NA |
| 4.066 | DCM04 | M | 52 | FDCM IHD | PKP2 | I531S | NA | NA | NA |
| S99 1036 | DCM05 | NA | NA | DCM | NA | NA | NA | NA | NA |
| TB11.1626 | DCM06 | M | 45 | DCM | NA | NA | White british | No | 15 |
| TB11.1730 | DCM07 | M | 41 | DCM | NA | NA | White british | No | 8 |
| TB12.0754 | DCM08 | M | 43 | FDCM | NA | NA | Asian british Indian | No | NA |
| TB12.1094 | DCM09 | M | 29 | DCM | NA | NA | White british | No | NA |
| TB13.0359 | DCM10 | M | 24 | DCM | NA | NA | White british | No | NA |
| TB13.0992 | DCM11 | M | 25 | DCM | NA | NA | White British | No | NA |
| TB13.1292 | DCM12 | M | 50 | DCM, NYHA calss III heart failure | NA | NA | Whit British | No | NA |
| TB13.2060 | DCM13 | M | 42 | DCM | NA | NA | White British | No | NA |
| TB13.2131 | DCM14 | M | 64 | DCM with biventricular dysfunction | NA | NA | NA | No | NA |
| TB13.2163 | DCM15 | F | 53 | Restrictive Cardiomyop athy | NA | NA | White British | No | NA |
| TB13.2277 | DCM16 | M | 37 | DCM | NA | NA | NA | NA | NA |
| TB14.0606 | DCM17 | M | 65 | DCM with RV ischaemia | NA | NA | NA | No | NA |

| | | | | | | | | | |
|---|---|---|---|---|---|---|---|---|---|
| TB14.1985 | DCM18 | M | 34 | DCM | NA | NA | White british | No | NA |
| TB14.2015 | DCM19 | M | 44 | DCM | NA | NA | White british | Type II | NA |
| TB15.0559 | DCM20 | F | 31 | DCM | NA | NA | Black british carribean | Type II | NA |
| TB16.0638 | DCM21 | M | 52 | DCM | NA | NA | NA | No | NA |

**Supplemental Table 4.** Non-failing controls used in this investigation

| Random ID | Assigned ID | Source |
|---|---|---|
| S99 577 | NF01 | unused donor |
| Ch1 5.089 | NF02 | unused donor |
| Ch1 5.138 | NF03 | unused donor |
| LV 7.004 | NF04 | unused donor |
| LV 7.012 | NF05 | unused donor |
| LV 7.060 | NF06 | unused donor |
| LV. 7.072 | NF07 | unused donor |
| LV 8.010 | NF08 | unused donor |
| LV NB 5.048 | NF09 | unused donor |
| LV NE 5.086 | NF10 | unused donor |
| S99 1309 | NF11 | unused donor |

**Supplemental figures**

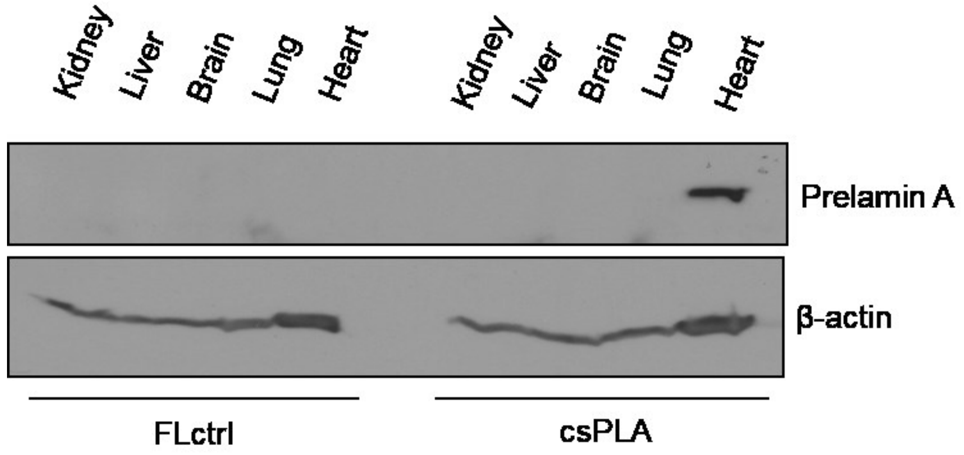


**Supplemental Figure 1. Prelamin A expression in csPLA-Tg mice was restricted to the heart.**

Western blotting of mouse tissues showed that prelamin A was expressed specifically in heart tissue.

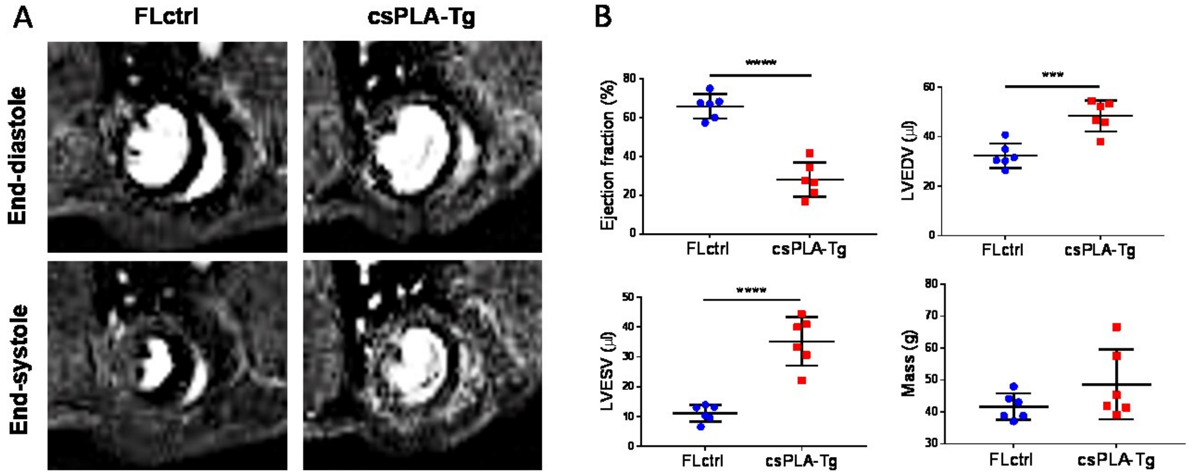


**Supplemental figure 2. Cardiac magnetic resonance imaging confirmed impairment in cardiac function of csPLA-Tg myocardium at four weeks.** A. Representative MR images of myocardium in end systole and end diastole. B. decrease in ejection fraction alongside increases in left ventricle end diastolic and systolic volume indicated dilated cardiomyopathy was the outcome of prelamin A accumulation. Mass was statistically unchanged but with increased variation and concurs with post-mortem heart weight measurements. Values are mean ± SD ***P<0.001, ****P<0.0001.

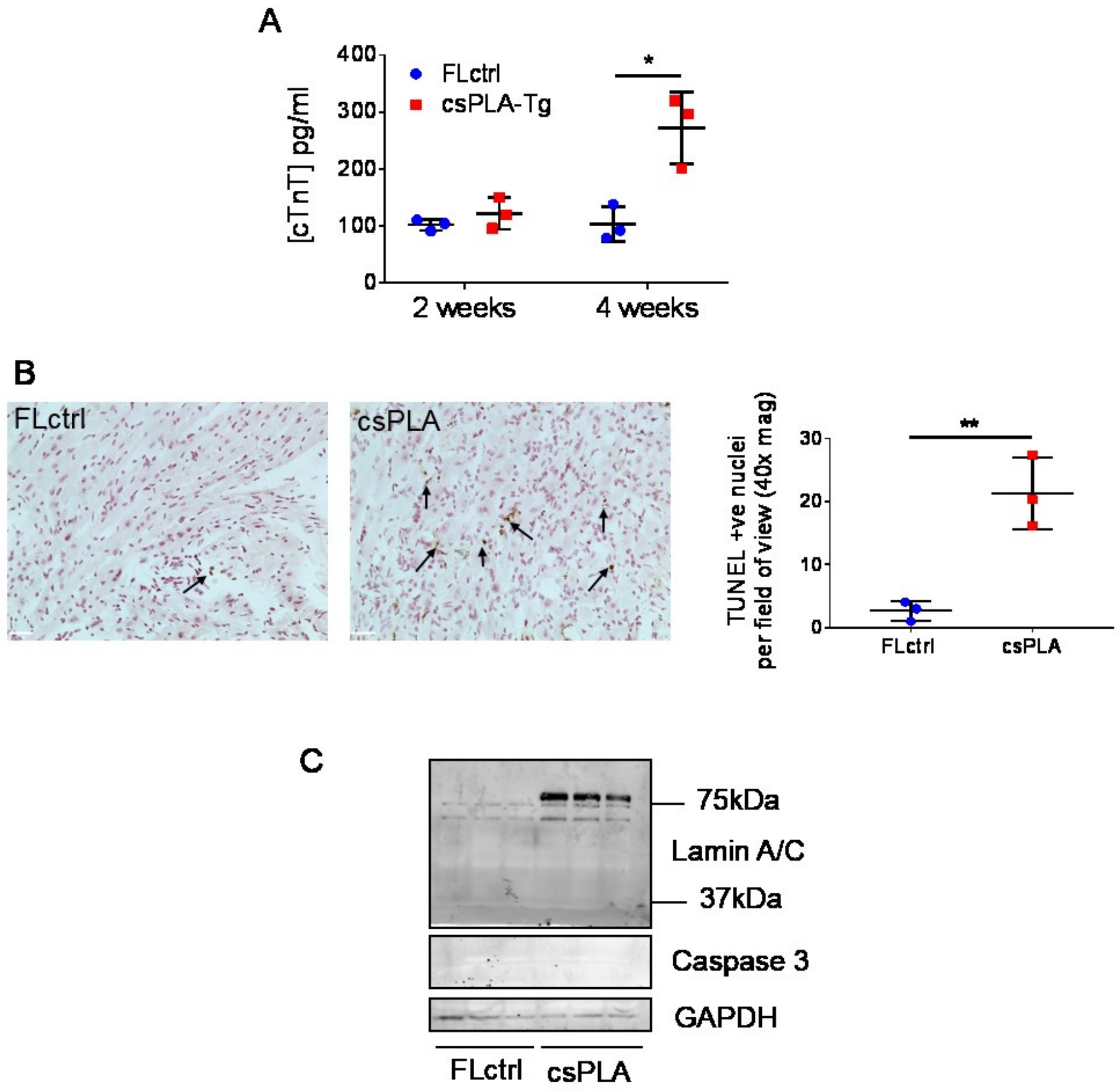


**Supplemental figure 3. Cell death observed in four week csPLA-Tg mice was not due to apoptosis. A.** Blood plasma subjected to ELISA for cardiac TNT showed elevated cardiac damage at in four week old mice**B.** TUNEL staining was performed on four week csPLA-Tg heart sections and image quantification showed a significant increase in TUNEL positive nuclei in csPLA-Tg heart sections at four weeks. **P<0.01. N=3/group **C.** Western blotting showed that cleavage of caspase 3 and lamins A/C, which occurs at ~37 kDa, which occur during apoptosis, did not occur in four week csPLA-Tg myocardium. Values are means ± SDs. *P<0.05, **P<0.01. N=3/group.

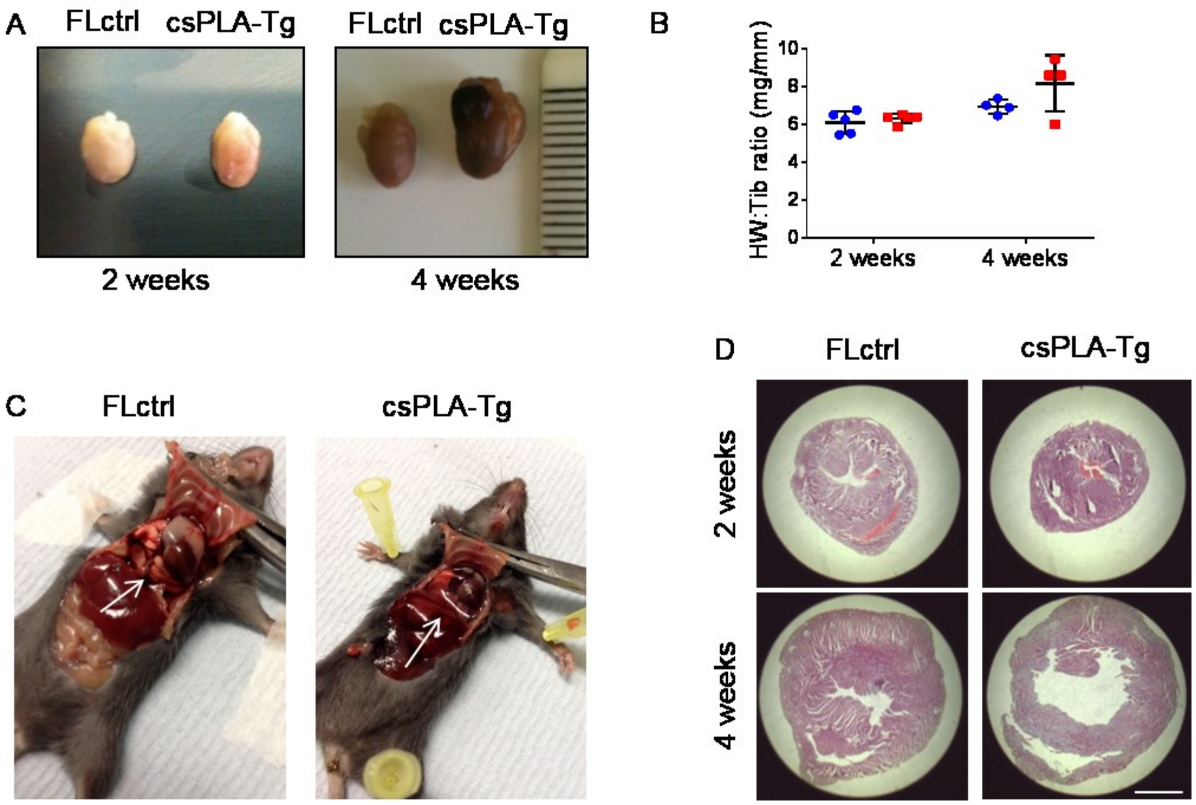


**Supplemental figure 4. csPLA-Tg mice hearts are dilated with evidence of heart failure. A.** At four weeks csPLA-Tg hearts were enlarged. **B.** No significant difference in heart weight to tibia length ratio was observed. Values are means ± SDs. N=3/group. **C.** Dissection of chest cavities showing transudative pleural effusions in csPLA-Tg mice. **D**. Histological assessment shows that heart chambers of four week csPLA-Tg mice were dilated. Scale = 2mm.

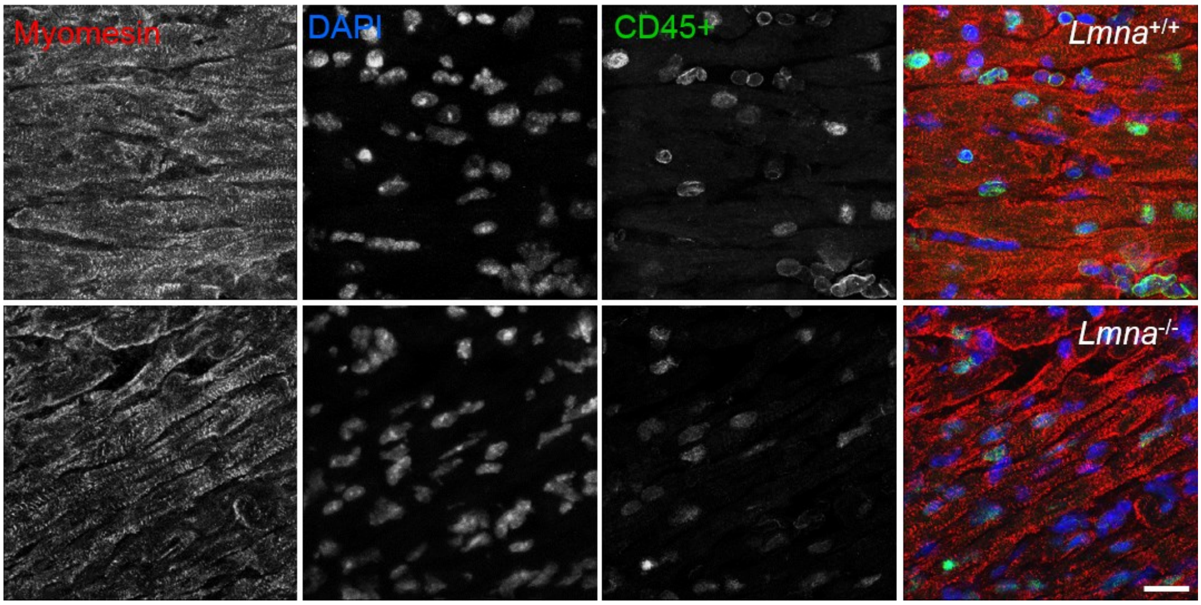


**Supplemental figure 5.** ***Lmna$^{-/-}$*** **mouse myocardium does not exhibit signs of inflammation.**

CD45+ immunostaining of *Lmna$^{-/-}$* myocardium showed no evidence of infiltration by leukocyte populations.
